# *pi_tailtrack*: A compact, inexpensive, and open-source behaviour-tracking system for head-restrained zebrafish

**DOI:** 10.1101/2023.06.01.543206

**Authors:** Owen Randlett

**Affiliations:** Laboratoire MeLiS, UCBL - CNRS UMR5284 - Inserm U1314, Institut NeuroMyoGène, Faculté de Médecine et de Pharmacie, 8 avenue Rockefeller, 69008, Lyon, France

## Abstract

Quantifying animal behavior during microscopy is crucial to associate optically recorded neural activity with behavioural outputs and states. Here I describe an imaging and tracking system for head-restrained larval zebrafish compatible with functional microscopy. This system is based on the Raspberry Pi computer, Pi NoIR camera, and open-source software for the real-time tail segmentation and skeletonization of the zebrafish tail at over 100hz. This allows for precise and long-term analyses of swimming behaviour, that can be related to functional signals recorded in individual neurons. This system offers a simple but performant solution for quantifying the behavior of head-restrained larval zebrafish, which can be built for 340€.

## Introduction

A chief application of the larval zebrafish for neuroscience is to image the activity of neurons in the intact and behaving animal using microscopy. This is facilitated by its translucent and small brain, measuring approximately 0.1 mm^3^. By expressing genetically encoded indicators, such as the GCaMP Ca^2+^ sensors (***Akerboom et al., 2012***; ***Chen et al., 2013***), signals related to the activity of practically any or all neurons can be recorded from the larval zebrafish brain (***Ahrens et al., 2012***; ***Portugues et al., 2014***).

Ca^2+^ imaging can be performed with standard microscopes, but such systems are not equipped for monitoring the behaviour of the animal. Therefore, any analyses directly relating neural activity to behaviour will require the integration of a behavioural recording apparatus. Behavioural recording is typically done in the context of custom-built microscopes, which can be designed explicitly with this behaviour-monitoring goal in mind. However, many groups (including my own) have neither the financial nor technical means to implement such a complete system. We rely on microscope equipment in a shared core facility. Such microscopes generally cannot be substantially or permanently modified, and often present physical and optical constraints that make installing a behaviour imaging system challenging.

Here I present a solution for this problem based on the Raspberry Pi computer, that I call *pi_tailtrack*. The system includes illumination, camera, computer and software, yielding a complete setup that is compact, inexpensive, and self-contained. The *pi_tailtrack* system can reliably track larval zebrafish behaviour in real-time at over 100hz while performing functional imaging experiments.

## Materials and Methods

### Animal Ethics Statement

Adult zebrafish used to generate larvae were housed in accordance with PRCI facility approved by the animal welfare committee (comité d’éthique en expérimentation animale de la Région Rhône-Alpes: CECCAPP, Agreement # C693870602). Behaviour and microscopy experiments were performed at the 5dpf stage, and are thus not subject to ethical review, but these procedures do not harm the larvae.

### Animals

All experiments were performed on larval zebrafish at 5 days post fertilization (dpf), raised at a density of ≈1 larvae/mL of E3 media in a 14:10h light/dark cycle at 28-29°C. Adult zebrafish were housed, cared for, and bred at the Lyon PRECI zebrafish facility. *mitfa*/Nacre mutant animals (ZDB-ALT-990423-22) were used to prevent pigmentation. Larval zebrafish were mounted and head restrained for 2-photon imaging and behavioural analysis by placing them in a very small drop of E3 in the lid of a 35mm petri dish (Greiner bio-one, 627102). Molten (≈42°C) 2% low melting point agarose (Sigma A9414) in E3 Medium was added to the dish in an approximately 10mm-diameter droplet around the fish, and the zebrafish was repositioned within the solidifying agarose using a gel-loading pipette tip, such that it was oriented symmetrically for imaging with the dorsal surface of the head at the surface of the agarose. After the agarose had solidified (≈10 minutes), E3 was added to the dish, and then the agarose around the tail was cut away. This was done using a scalpel in two strokes emanating laterally from just below the swim bladder (illustrated in ***Figure 3***A). It is critical to not scratch the dish in the vicinity of the freed tail, which can interfere with tail-tracking.

### Hardware

I used a Raspberry Pi 4 Model B Rev 1.4 computer, running Raspbian GNU/Linux 11 (bullseye). ***Table 1*** contains the details of the hardware components that I used, their approximate price, and an option for supplier (keeping in mind that these later two are subject to change and will rapidly become inaccurate).

**Table 1.**
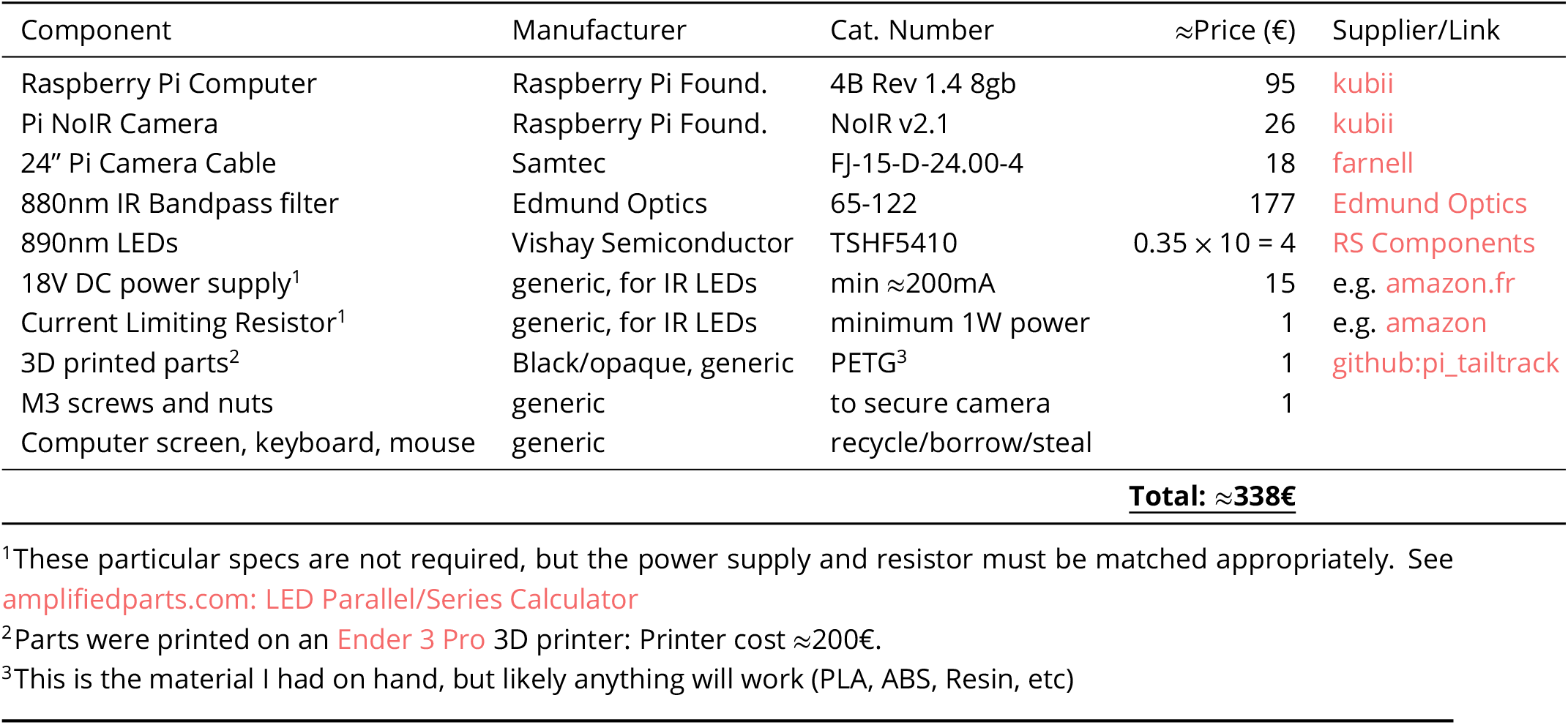
Bill of Materials.

The short 2 cm focal distance between the animal and the camera allowed for a compact and direct imaging setup, where the camera is mounted directly below the larva (***Figure 1***). This avoids the need for any mirrors, and frees the space above the animal for the microscope objective, and any stimulus apparatus necessary. In our case we use red LEDs to provide visual stimuli to the larvae (***Lamiré et al., 2022***).

**Figure 1.**
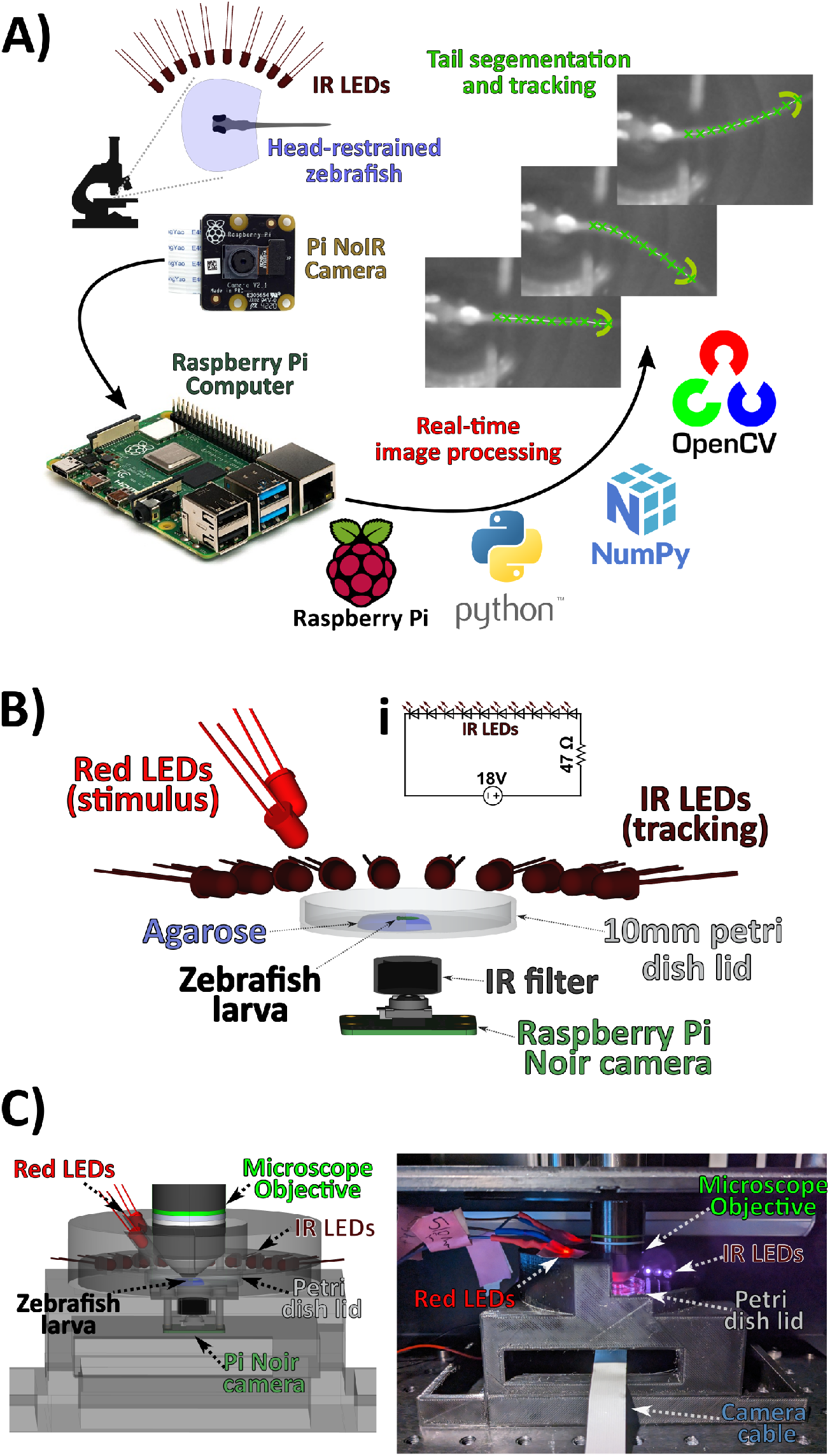
*pi_tailtrack* apparatus. A) The zebrafish larvae being imaged under the microscope is illuminated with infra-red (IR) LEDs, and imaged with the IR-sensitive Raspberry Pi NoIR Camera. Image acquisition and processing is done with a Raspberry Pi Computer and open-source Python packages. The zebrafish tail is identified and segmented in real-time as a sequence of 10 tail segments (green X’s). B) Rendering of the main components of the apparatus. IR leds illuminate the zebrafish larvae that is head-restrained in agarose in a 35mm diameter petri dish lid. An IR filter blocks the visible stimulus lights (Red LEDs), and the microscope laser from reaching the Raspberry Pi NoIR camera suspended below the fish. **(i)** Wiring diagram for powering the IR LEDs. C) Rendering including the the 3D printed mount and microscope objective. D) Annotated photograph of the *pi_tailtrack* apparatus.

To illuminate the larvae and visualize the tail, I used 890nm IR LEDs. Using the IR LEDs as an oblique illumination source generated a nicely resolved image of the *mitfa* mutant zebrafish tail that was sufficient for reliable identification and tracking (***Figure 2***). IR LEDs were wired in a simple circuit, with 10 LEDs in a series, powered by a 18V DC power supply and a 47ohm current limiting resistor (***Figure 1***Bi). Using these exact Voltage/Resistance configurations is not important, provided a relevant power supply and resistor are chosen to match the LED characteristics (forward voltage =1.4V, current = 100mA, for our 890nm LEDs: see for example amplifiedparts.com: LED Parallel/Series Calculator).

**Figure 2.**
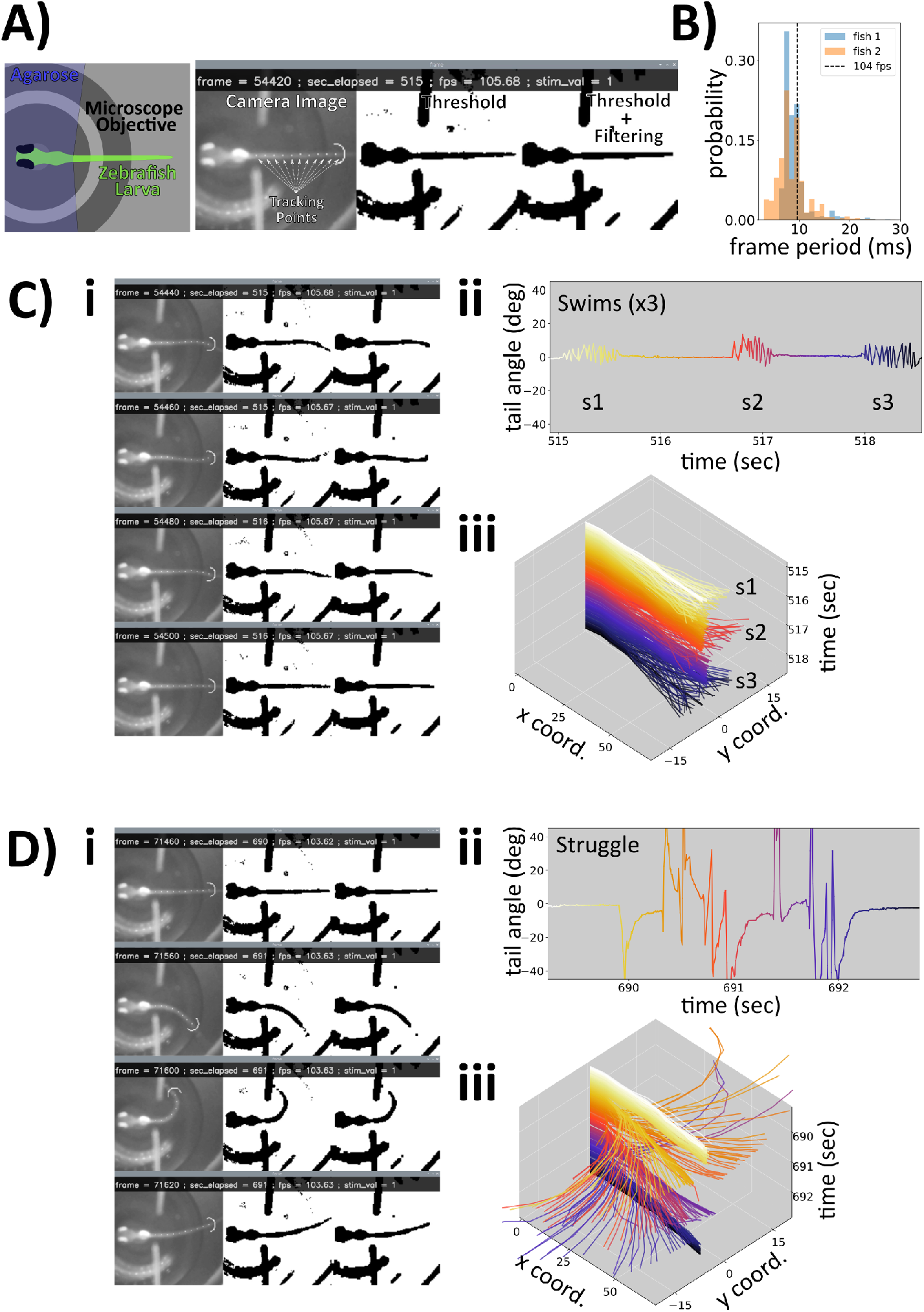
Larval zebrafish trail tracking examples. A) Screenshot of a single frame of a tracking image, showing the image from the camera (“Camera Image”) with the resultant tracking points overlayed as white dots. The final tracking point is shown as a white semicircle, which is used in the coordinate search algorithm. “Threshold” shows the result of the Adaptive Thresholding operation, and “Threshold + Filtering” the result of the morphological Opening and Closing operations. Displayed along the top are the: frame (current frame number of the experiment), sec_elapsed (number of seconds elapsed in the experiment), fps (current frame rate, frames per second), stim_val (the current value read on the stimulus recording pin: GPIO Pin 4). A schematic of the image field, depicting the agarose mounting medium, the position of the zebrafish, and the microscope objective visible in the background is shown in the left panel. B)Probability density distribution of individual frame periods from two representative experiments. C)Example frames during a swimming event. ii) Tail angle deflections during 3 distinct swim events. iii) 3D plot of tail coordinates through the same time period as (ii), drawn in the same time color code. D)Same as (C), but for a period in which the larvae executes a struggle/escape maneuver and associated high amplitude tail deflections. **Figure 2—video 1**. Screen recording of the tail tracking example, download

**Figure 3.**
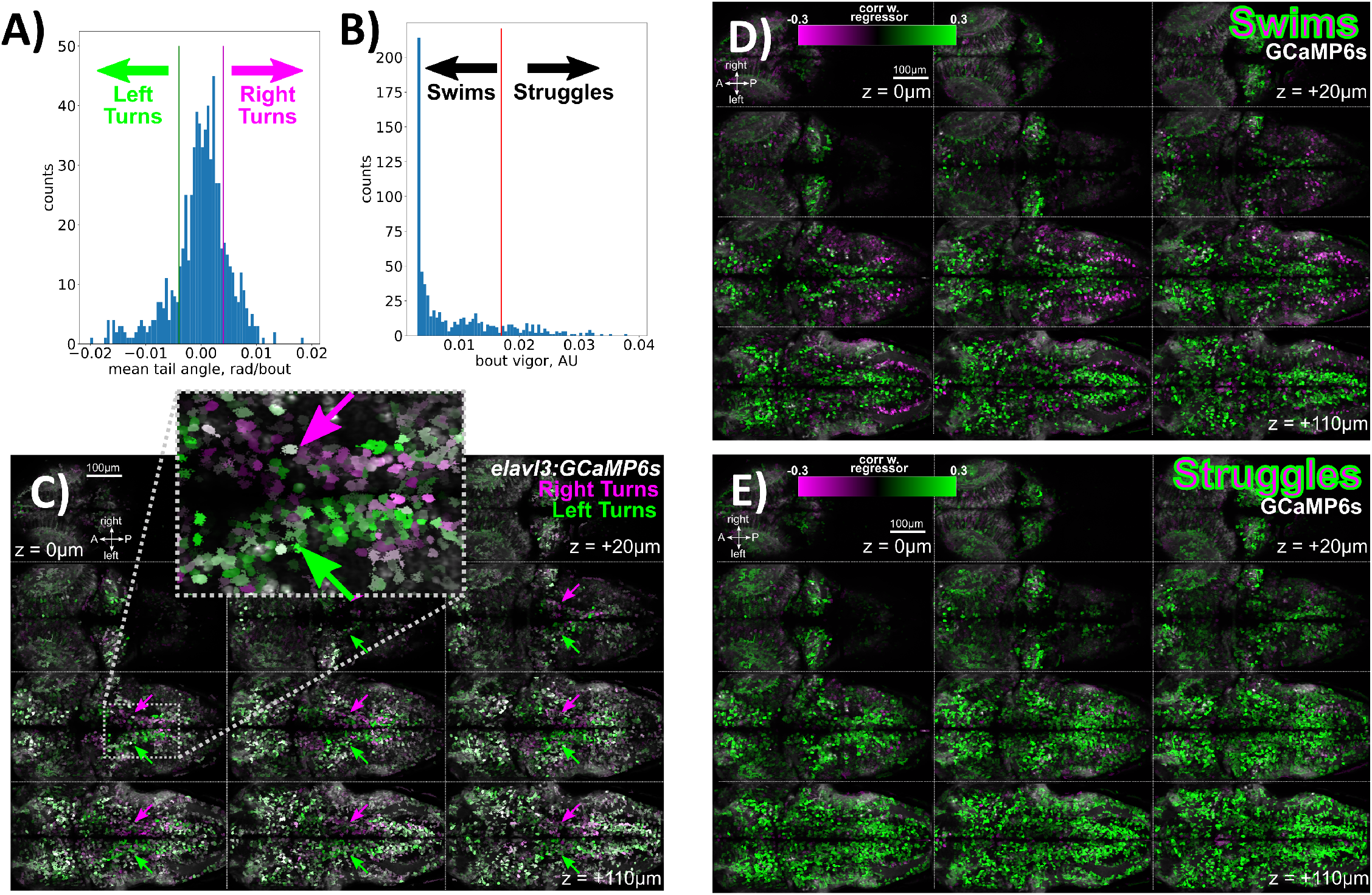
Identification of behaviour-associated neurons in a larval zebrafish brain via 2-photon Ca^2+^ imaging. A) Histogram for the mean tail angle during individual movement bouts for a single larvae over an 80 minute imaging session. Bouts are classified as left or right turns based on a threshold value of 0.004 radians/bout. B) Histogram for the bout vigor, quantified using a rolling standard deviation of absolute tail angle. Movements are classified as “swims” or “struggles” based on a threshold value of 0.017 (AU: arbitrary units). C)Tuning of Ca^2+^ traces in ROIs to turns to the left (green) or right (magenta), as classified in (A). Images are the Pearson correlation coefficient to each behavioral regressor (left or right turns), scaled from 0.0 to 0.3. *Tg2(elavl3:GCaMP6s)* expression pattern is shown in grey. Arrows highlight the Anterior Rhombencaphalic Turning Region (ARTR): with isilateral tuning to turning direction. A = Anterior, P = Posterior **D, E)** Tuning of neurons to swims **(D)**, and struggles **(E)**, as classified in (B).

We used an 880 nm bandpass filter in front of the Raspberry Pi NoIR camera module to selectively pass the IR LED light. This filter is essential to block the intense microscope laser light, which will obscure the image of the fish by saturating (and likely damaging) the camera sensor. Notably, this filter it is the most expensive part in the setup, costing more than the computer and camera, combined (***Table 1***). With our typical 2-photon GFP/GCaMP imaging settings and the laser tuned to 930nm, laser light is not visible in the camera image. Using such a bandpass filter in the 880 nm range should allow this system to be compatible with many other imaging modalities (c that the excitation wavelengths are not in the ≈870-900nm range, and the microscope system effectively filters out the 890nm light from the LEDs. If necessary, these wavelength characteristics can be adapted using different LED and filter components.

To house the system components I used a 3D printed mount (***Figure 1***C,D). This was designed using FreeCAD (freecad.org, FreeCAD file), and 3D printed in black PETG using and Creality Ender 3 Pro 3D printer. It consists of the main body shape that holds the the camera, IR filter, red stimulus LEDs above the fish, and IR LEDs in the oblique illumination configuration (Main Shape). An insert is placed into the depression above the IR filter, forming the platform onto which the zebrafish dish is placed (Depression Insert). The final 3D printed component is a semicircular shape that completes the encirclement of the objective, and helps minimize light scattering (***Figure 2***Bi, Semicircle STL file).

I would note that I built up the size of the platform of the mount to match with the relatively spacious configuration of the microscope I was using (***Figure 1***D). A much more compact configuration is possible, since we only require ≈26 mm of clearance from the fish to the bottom of the ≈6 mm thick camera. The base design could be easily adapted to match different microscope stage configurations. For example the entire system could be inverted to accommodate an inverted microscope to image ventral structures during behaviour, such as the the lateral line ganglia or the heart. Or, if stimuli need to be bottom-projected, a small 45-degree hot mirror could be used to divert the image to the camera and free the space directly beneath the animal for stimuli.

### Tail Tracking Approach

Software was written in Python, using the *picamera* library for camera control (***Raspberry Pi Foundation***, ***2023***). Trail tracking was performed using *OpenCV* (cv2 version 4.5.5) (***Bradski***, ***2000***), and *Numpy* (version 1.19.5) (***Harris et al., 2020***). All code is provided in the file record_tail.py.

Image frames are acquired directly from the camera buffer as an 8-bit Numpy array, and thresholded using Adaptive Thresholding (*cv2*.*adaptiveThreshold*) to identify bright objects in the image (***Figure 2***A, “Threshold”), using a threshold of −10 and a 33 pixel neighborhood. This binary image is then filtered using a morphological Opening and Closing operation (*cv2*.*morphologyEx*). This combination generally results in a nicely segmented fish blob in the final binary image (***Figure 2***A, “Threshold + Filtering”). Thresholding and filtering parameters can be adjusted in real-time using the w/s and a/d keys. However, this method identifies all large bright objects in the image, including borders of the agarose block and reflections on the microscope objective, and therefore we need a method to identify the fish object among these various segmented blobs.

The fish object is identified with a pre-defined coordinate that acts as the first tracking point of the fish. The fish object is then skeletonized into up to 10 tail segements (***Figure 2***A, ‘Tracking Pts’), which can be used to reconstruct the posture of the tail to identify swimming events (***Figure 2***C,D). To do this skeletonization, the tracking points are iteratively identified based on the intersection of a semicircle and the fish object, offset 7 pixels (0.19mm) from the previous tracking point, and oriented in the direction of the previous segment (similar to ***Štih et al. (2019***); ***Randlett et al. (2019***)). For the first search, this direction is toward the right of the image. Therefore, this strategy relies on the zebrafish larvae being oriented with its tail pointed towards the right, and being placed in the same position such that the exit point of the tail from the agarose block intersects with the first tracking point. The starting coordinate for the tail tracking can be adjusted using the arrow keys. It also requires that no other bright objects intersect with the fish object after binarization. Therefore, it is critical to avoid distracting objects in the imaging scene, such as scratches in the dish or stray pieces of agarose.

### Tracking data format

The tail tracking data are saved in a comma-separated text file *‘*_coords*.*txt’*, the 10 pairs of “X” and “Y” coordinates for each tail point are saved as rows, and thus there are two rows with 10 columns for every tracked frame. ‘NaN’ values represent instances where a tail point is not identified.

The timing of the data is saved in a separate text file *‘*_tstamps*.*txt’*, which also has two rows for each frame. The first value is the “timestamp” reflecting the time elapsed since the beginning of the tracking experiment. This is used to relate the tail posture and behavioural events to specific points in time. This is important for experiments in which precise timing of behavioural events is important, because the frame rate is not fixed and can fluctuate during the experiment (see above). However, it is important to note that the timestamp recorded is based on the time at which the frame is received from the camera buffer, which may lag from the time at which it was actually acquired by the camera. This could be problematic if millisecond-level precision on behavioural timing is critical, for example if differentiating between Short- and Long-Latency acoustic stimulus responses (***Burgess and Granato***, ***2007***).

The second value in the *‘*_tstamps*.*txt’* file is the value recorded on one of the GPIO pin 4 of the Raspberry Pi. This value will read either “low”=0 for a voltage less than 1.8V, or “high”=1 for 1.8-3.3V. I use these recordings to synchronize the behavioural recordings with the frames recorded on the microscope. In our typical setup we are using an analog output pin from the DAQ board on the microscope to control the red stimulis LEDs (***Figure 1***B), and we also connect this output of the DAQ board to GPIO pin 4 on the microscope. In this way, we can synchronize the stimuli, microscope imaging frames, and the behavioural recordings.

These datasets can be read into python for analysis using, for example:

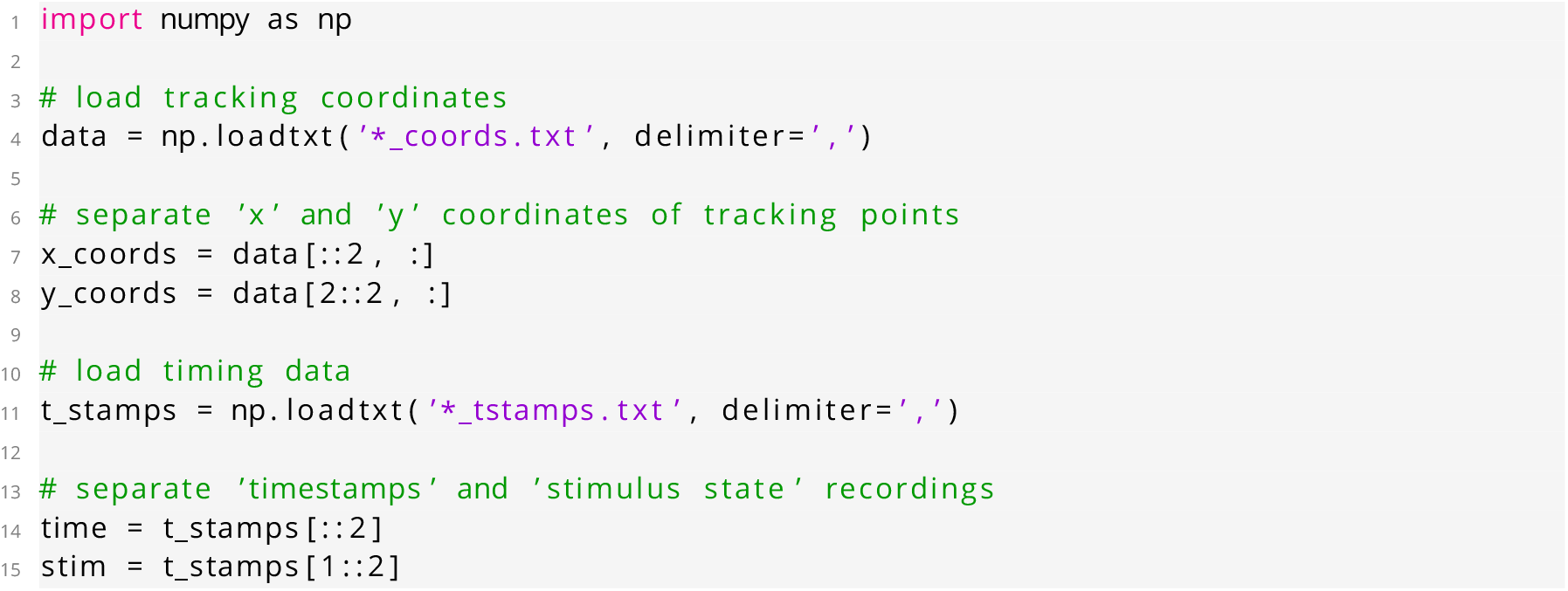

### Ca^2+^ imaging and analysis

2-photon Ca2+ imaging was performed and analyzed as described in (***Lamiré et al., 2022***). Briefly, a 5dpf *Tg2(elavl3:GCaMP6s)* (ZDB-ALT-180502-2, ***Dunn et al. (2016***)) larva was imaged using a 20x 1.0NA water dipping objective (Olympus) on a Bruker Ultima microscope at the CIQLE imaging platform (Lyon, LYMIC). Frames were acquired using a resonant scanner over a rectangular region of 1024×512 pixels (0.6µm x/y resolution) and piezo objective to scan 12 planes separated at 10µm steps, with a repeat rate of 1.98 hz. The position of the functional imaging stack within the brain was stabilized in x/y/z “online” during acquisition using Bruker’s PrairieLink API and Python (brukerPL_stable_tseries.py). The central imaging plane was compared to a high-quality anatomical stack acquired before functional imaging using the *registration*.*phase_cross_correlation* function from the *scikit-image* package (***Van der Walt et al., 2014***).

ROIs were identified and fluorescence timeseries extracted using suite2p (***Pachitariu et al., 2016***). The zebrafish was stimulated with 60 “dark flash” stimuli at 60 second ISI (***Lamiré et al., 2022***), though responses to these stimuli were not incorporated into the analyses presented here, other than to synchronize the behavioural tracking with the microscope acquisition timing.

To identify neurons tuned to turning direction (***Figure 3***C), the fluorescence trace from each ROI was compared to vectors derived from the *pi_tailtrack* recordings of the tail reflecting leftward or rightward turns, respectively. These “behaviour state” vectors were convolved with the GCaMP response kernel to generate “regressors” reflecting the predicted Ca^2+^ response in neurons that are activated during the relevant behavioural state (as in ***Miri et al. (2011***)). Tuning images were then generated reflecting the Pearson correlation coefficient between the z-scored fluorescence trace of the ROI and the relevant regressor. Images output from the analysis were adjusted for brightness/contrast and LUT using FIJI/ImageJ (***Schindelin et al., 2012***). This same approach was used to identify the relationship between ROIs and “Swim” and “Struggle” motor events (***Figure 3***D,E).

## Results and Discussion

### Design goals

I wanted to track the swimming behaviour of head-restrained larval zebrafish while performing Ca^2+^imaging. There are many ways that this might be accomplished, but I wanted a system that was:

1. Able to identify and characterize individual swimming events while we are imaging the brain using 2-photon microscopy.
2. Compact and self contained, so that it can be easily and rapidly installed and removed for our imaging sessions on a shared microscope.
3. Made using low-cost and open source hardware and software to facilitate re-use in other contexts, and because I am a cheap *financially responsible* researcher.

### Using a Raspberry Pi camera to image the larval zebrafish tail

The Raspberry Pi is a very inexpensive, credit-card-sized computer that plugs into a standard monitor, keyboard, and mouse. The Raspberry Pi’s open-source nature and large user community, and its ability to control and interface with a variety of devices and sensors make it a powerful and accessible platform for developing and sharing custom neuroscience and behavioural research tools. Indeed many such systems have been developed in recent years based around the Raspberry Pi and the Pi Camera, and especially the IR-sensitive Pi NoIR camera, as an acquisition device (***Geissmann et al., 2017***; ***Maia Chagas et al., 2017***; ***Saunders et al., 2019***; ***Tadres and Louis***, ***2020***; ***Broussard et al., 2022***).

However, obtaining sufficient resolution and contrast to resolve the larval zebrafish tail is challenging since the tail is very narrow (≈0.25mm diameter), and nearly transparent. This is especially true in de-pigmented animals that are generally used for brain imaging due to their lack of melanin pigment over the brain (e.g. *mitfa*/Nacre mutants, or larvae treated with N-Phenylthiourea). This also removes melanin pigment from the tail, increasing its transparency and making it harder to image and track. Thus, it was not clear if the 26€ Pi NoIR Camera would be up to this task.

The stock lens configuration on the Pi Camera is also not designed for macro photography, and has a minimum focus distance of 50cm. But, extension tubes are a well-known macro-photography hack that work by increasing the distance between the lens and the camera (***Wikipedia contributors***, ***2022***). Increasing this distance acts to decrease the focus distance of the optical system, increasing the maximal magnification. By unscrewing the lens of the Pi NoIR camera until just before it falls off, it is possible to focus on objects at a 2 cm distance, allowing for sufficient magnification to observe the and track the tail of *mitfa* mutant zebrafish (***Figure 1, Figure 2***).

A second challenge is that larval zebrafish move their tails very rapidly, with a tail-beat frequency of between 20-40 hz for normal swimming, which can increase to 100 hz during burst/escape swimming (***Budick and O’Malley***, ***2000***; ***Muller***, ***2004***; ***Severi et al., 2014***). The V2.1 camera documentation indicates maximum frame rate of 30hz, which is insufficient for imaging tail dynamics. However, by adopting a cropped sensor configuration, and by omitting the JPG compression step in image processing, the camera can be pushed to image at up to 1000hz (***Elder***, ***2019***). I adopted a configuration where I imaged with a fixed gain/ISO of 800 in auto-exposure mode, and with a cropped sensor of 128×128 pixels covering 3.5×3.5 mm field of view. This gives sufficient spatial resolution to observe and track the tail of the fish (27 µm/px), and most importantly, minimal CPU load. This frees the limited CPU resources on the Raspberry Pi to be used for real-time image processing and tail tracking.

### Tail tracking

Tracking objects in images and videos has undergone a revolution with deep learning and neural network frameworks, where the tracking and reconstruction of complex animal postures is possible after training networks on only a few example images (***Mathis et al., 2018***; ***Pereira et al., 2022***). However, such approaches are computationally intensive and generally require dedicated and GPU hardware beyond the capabilities of the standard Raspberry Pi, making them incompatible with our project goals. In contexts where the image background is predictable and stable, classical computer vision methods like background subtraction, filtering and thresholding may still be preferable to network-based object identification, especially when speed or computational resources are priorities (***Mirat et al., 2013***; ***Štih et al., 2019***; ***Zhu et al., 2023***). Here I have used the *Numpy* (***Harris et al., 2020***) and *OpenCV* (***Bradski***, ***2000***) libraries to handle the image data and computer vision tasks (***Figure 1***).

I used a computationally lean segmentation and skeletonization strategy (see Materials and Methods) to segment the tail into 10 segments (***Figure 2***), which takes less than 10 ms on the Raspberry Pi CPU. The imaging frame rate when using the *picamera* python package will adjust based on the throughput of the analysis system, which can change with the complexity of the binary images that are processed or external CPU demands, but runs at approximately 104 fps (***Figure 2***B). This is sufficient to clearly distinguish different types of movement events, such as “swims” from “struggles” ***Figure 2***C vs. D), and where individual tail beats during swimming events are resolvable. However, this will not be true during rapid/burst swimming, in which tail-beat frequency will exceed our frame rate. If such temporal resolution is required our setup will be insufficient, and we will will only reliably track tail half-beat frequencies of ≤50hz. Therefore, this system is not capable of comprehensive behavioural characterization, but can be used to identify different types of swim events.

During the experiment the software provides a visual display, as is shown in the screenshots in (***Figure 2***), and screen capture video (***Figure 2***–***video 1***). Results of the thresholding, filtering, and skeleton tracking are visible and updated in real-time. This can be used to optimize the position of the zebrafish, the Adaptive Thresholding parameters (neighborhood, threshold) using the *‘w/a/s/d’* keys, and the position of the first tracking point using the arrow keys.

### Behavioural analysis of Ca^2+^ imaging data

To test the performance of the *pi_tailtrack* system, I analyzed Ca^2+^ imaging data from an 80 minute-long volumetric recording covering a large proportion of the brain (as in ***Lamiré et al. (2022***)). To identify neurons tuned to behavioural parameters I used “regressors” derived from the *pi_tailtrack* recordings reflecting different motor states convolved with the GCaMP response kernel (as in ***Miri et al. (2011***)). Zebrafish swim bouts can be classified as either forward swims or turns, and an area within the anterior hindbrain is associated with turning direction. This area is known as the Anterior Rhombencephalic Turning Region (ARTR: ***Dunn et al. (2016***), also called the HBO: ***Ahrens et al. (2013***); ***Wolf et al. (2017***)), and shows a conspicuous activity pattern with stripes of neurons tuned to the ipsilateral turning direction. By looking at correlations to regressors reflecting right and left turns, I identified these stripes of neurons in the ARTR-region, indicating that I can successfully identify the ARTR using *pi_tailtrack* (***Figure 3***A,C). A similar analysis looking at “swims” vs “struggles”, with “struggles” reflecting high-amplitude tail flicking events (***Figure 2***D, ***Figure 3***B), identified differential neuronal activation in the context of these two movement categories (***Figure 3***D,E), with the presence of lateral hindbrain populations of neurons that were negatively correlated with “swims”, and a broader and more positively correlated population with “struggles”.

### Future developments

Here I have used the *pi_tailtrack* system to simply record the behaviour of the animal independent of the microscopy or any stimulus delivery. Therefore, the timing of microscope image acquisition is controlled by the microscope computer and is independent of *pi_tailtrack*. These separate experimental clocks (microscope frames vs Pi Camera frames) must be synchronized, and in our case I have used the GPIO input pin on the Raspberry Pi to record the timing of the stimuli delivered by the microscope relative to the Pi Camera frames. An alternative solution would be to use the Raspberry Pi to deliver the stimuli, perhaps by integrating a video projector system to allow for the delivery of arbitrary and complex visual stimuli. This would also open up possibilities for performing “virtual reality” experiments, where the behaviour of the animal dictates the stimulus in closed-loop. In some microscope systems it should also be possible to use the Raspberry Pi GPIO to trigger microscope acquisitions. This may be preferable if the synchronization between imaging and behaviour frames is critical.

It is also important to note that hardware in this micro-computer/Raspberry Pi space is rapidly evolving. Indeed, a new suite of Raspberry Pi V3 Cameras have just been released, offering increased resolution, dynamic range, and frame rate. Using these cameras, we may be able to increase the frame rate of tracking into the multiple-hundreds of hz, which would allow us to more reliably resolve individual tail half-beats. The Raspberry Pi “Global Shutter” Camera has also recently been released, which is likely also going to be very interesting for behavioural neuroscience, as the use of a global shutter avoids rolling shutter artifacts that distort images along the frame during rapid motion. The software introduced here could be further optimized for speed/framerate, for example by moving to a multi-threaded architecture to distribute the image acquisition and tracking computations (***Zhu et al., 2023***; ***Randlett et al., 2019***), using a compiled language (e.g. C/C++ or Julia), or perhaps by moving image processing onto the Raspberry Pi GPU.

## Conclusion

Here I described our system for tracking the tail of the larval zebrafish during microscopy. Many of the practical considerations of this setup may be specific to our application, and therefore may need modification for use in other experiments in other labs. However, I feel that the core and simple idea of using an IR-sensitive Raspberry Pi Camera, a simple Python script, coupled with IR LEDs and and IR filter, provides an approachable and flexible solution that may be widely useful for observing and tracking the behaviour of zebrafish (or perhaps other animals) while performing imaging experiments. This system’s attributes may also make it an ideal tool for community engagement activities such as school outreach programs. It could serve as a platform for learning about microelectronics, behavioural analyses, machine vision, and hardware design and construction.

## Funding

This work was supported by funding from the ATIP-Avenir program of the CNRS and Inserm, a Fondation Fyssen research grant, and the IDEX-Impulsion initiative of the University of Lyon.

## Data Availability

Software and analysis code is available here: https://github.com/owenrandlett/pi_tailtrack/. Datasets are available here: [pi_tailtrack datasets](https://www.dropbox.com/sh/dbjq2dud1ws1o2v/AACLamthISys8sUD1a5oRcR1a?dl=0).

## Notes

https://github.com/owenrandlett/pi_tailtrack

### Competing Interest Statement

The authors have declared no competing interest.

### Summary of Updates

Updated based on suggestion from reviewers. Consolidated original figure 1 and 2 into a single figure. Added some technical details.

